# MAPPING CHANGES OF MIRNA-MRNA NETWORKS IN *LEISHMANIA-INFECTED* MACROPHAGES PREDICTS REGULATORY MIRNA-TF LOOPS AS NOVEL TARGETS OF PARASITE IMMUNE SUBVERSION

**DOI:** 10.1101/2024.03.24.586456

**Authors:** Charfeddine Gharsallah, Hervé Lecoeur, Hugo Varet, Rachel Legendre, Odile Sismeiro, Jean-Yves Coppée, Caroline Proux, Elisabetta Scarfiello, Astrid Bruckmann, Gunter Meister, Eric Prina, Gerald F. Späth

## Abstract

MicroRNAs (miRNAs) are small non-coding RNAs that regulate gene expression at the post-transcriptional level and play a crucial role in numerous disease processes, including infections. Although intracellular microbial pathogens are known to modulate host cell gene expression to establish permissive conditions for infection, the specific role of host-encoded miRNAs underlying such subversion remains poorly understood. In this study, we employed the protozoan parasite *Leishmania amazonensis* as a model system to investigate how infection of macrophages modifies the host cell miRNA profile to evade antimicrobial functions and to establish permissive conditions for intracellular proliferation. Dual RNA-seq analyses using matched mRNA and miRNA-enriched samples from uninfected and *L. amazonensis*-infected bone marrow-derived macrophages (BMDMs) revealed 102 differentially expressed miRNAs (padj<0.05), with 18 miRNAs showing reduced and 84 miRNAs showing increased abundance in infected BMDMs. Mapping putative networks of miRNA-mRNA interactions based on the observed expression changes, combined with Gene Ontology enrichment analyses, allowed us to identify potential miRNA target genes involved in key biological processes and metabolic pathways that permit parasite intracellular survival and proliferation. Our analyses predict the existence of a large miRNA-mRNA network affecting the expression level of numerous transcription factors that indicates inhibition of the NF-κB-dependent inflammatory response or the promotion of cholesterol biosynthesis during infection. In particular, the over 10e3-fold increase in the abundance of mmu-miR-686 in infected BMDMs was correlated with a reduced abundance of putative target transcripts implicated in miRNA biogenesis itself, in RNA binding, and in regulation of apoptosis, such as *Caspase 12*, the mRNA decay activator protein *Zfp36l1* or *Leukemia Inhibitory Factor Receptor Alpha*. Likewise, the over 200-fold increase in abundance of mmu-miR-6546-3p was associated with a reduced abundance of putative target mRNAs implicated in cytokine-mediated signaling, positive regulation of apoptotic process and regulation of gene expression, affecting, for example, the *MADS box transcription enhancer factor 2*, the *transformation related protein 53 inducible nuclear protein 1*, or the *G protein-coupled receptor 35*. Interestingly, both miRNAs are predicted to simultaneously target 32 mRNAs that showed reduced abundance in infected BMDMs, including *Maturin Neural Progenitor Differentiation Regulator* (*Mturn*), a regulator of NF-κB transcription factor activity. In conclusion, our approach provides novel insight into molecular mechanisms that may govern macrophage subversion and intracellular *Leishmania* survival. Our results shed new light on the complex relationship among miRNAs, macrophage gene expression and *Leishmania* infection, proposing regulatory feed-forward loops (FFLs) and feedback loops (FBLs) between miRNAs and TFs as a novel target of *Leishmania* immune subversion. These findings open exciting new avenues for the development of intervention strategies aimed at disrupting such crucial interactions, for example using an anti-miR (antagomir) approach against mmu-miR-686 and mmu-miR-6546-3p.

## INTRODUCTION

Epigenetic regulation is a complex mechanism that fine-tunes eukaryotic gene expression. Among epigenetic factors, microRNAs (miRNAs) have emerged as key molecules, intervening at the post-transcriptional level by targeting specific messenger RNAs (mRNAs) to modulate gene expression. Their implications extend to a wide range of human diseases, encompassing areas as diverse as cancer and microbial infections (Atri et al., 2019; Zhang et al., 2019). The importance of miRNAs lies in their ability to act as subtle regulators, actively participating in the precise adjustment of protein levels in cells. This regulation is crucial for maintaining cellular homeostasis and for the adaptive response to environmental challenges (Sprenkle et al., 2023). However, the influence of miRNAs is not limited to normal gene regulation. Indeed, they play a leading role in the context of microbial infections, where pathogens - whether viral, bacterial, or eukaryotic - have developed sophisticated epigenetic mechanisms to disrupt gene expression within host cells, thus ensuring their survival and replication (Ahluwalia et al., 2017; Sun et al., 2017). One of the strategies favored by these pathogens is the modulation of the expression of host miRNAs.

*Leishmania* are protozoan parasites that infect mammalian macrophages, inducing a significant disruption of the functions of these cells, leading to severe immune pathologies known as “leishmaniases” (Conceicao-Silva and Morgado, 2019; McGwire and Satoskar, 2014). The consequences of this immune subversion are varied, ranging from self-healing cutaneous forms to potentially fatal visceral forms. However, despite the magnitude of the public health problems posed by leishmaniasis and the profound impact of *Leishmania* infections on the physiology of host cells, the precise mechanisms of this immune subversion remain incompletely understood.

Recently, epigenetic subversion has emerged as a major strategy that these parasites use to escape host defenses (Gutierrez Sanchez et al., 2023; Kamhawi and Serafim, 2020; Lecoeur et al., 2022a). First, *Leishmania* infection has been correlated to changes in host miRNA abundance, known to limit the expression of pro-inflammatory genes, thus favoring parasite survival (Ganguly et al., 2022; Lemaire et al., 2013; Mukherjee et al., 2015; Muxel et al., 2018; Nandan et al., 2021; Tiwari et al., 2017). Second, intracellular infection by *Leishmania* causes important changes in the macrophage DNA methylation pattern, affecting signaling pathways, oxidative phosphorylation, and cell adhesion (Marr et al., 2014). Finally, we have previously provided the first evidence that *Leishmania* also manipulates the histone code of the host cell both *in vitro* and *in vivo*, causing hypo-methylation and hypo-acetylation at promoters of pro-inflammatory genes that correlated with reduced gene expression (Lecoeur et al., 2020a).

Here, we performed a dual RNAseq analysis to simultaneously profile changes in mRNA and miRNA abundances in primary, bone marrow-derived macrophages infected with lesion-isolated *L. amazonensis* amastigotes (Fernandes et al., 2022; Lecoeur et al., 2022b; Lemaire et al., 2013; Martinez-Hernandez et al., 2021; Rashidi et al., 2021). Computational analysis of these data sets allowed us to map putative miRNA-mRNA regulatory networks that are predicted to control the observed inhibition of the pro-inflammatory TLR / NF-κB signalling pathway and increased cholesterol biosynthesis, both of which are prerequisites for intracellular parasite survival (Gutierrez Sanchez et al., 2023; Lecoeur et al., 2020a). Our findings open new perspectives for anti-parasitic intervention targeting the non-coding transcriptome of the host cell.

## MATERIAL AND METHODS

### Ethics statement

Six-week-old female C57BL/6JRj and Swiss nu/nu mice were purchased from Janvier (Saint Germain-sur-l’Arbresle, France). All animals were housed in A3 animal facilities according to the guidelines of Institut Pasteur and the “Comité d’Ethique pour l’Expérimentation Animale” (CEEA) and protocols were approved by the “Ministère de l’Enseignement Supérieur, Direction Générale pour la Recherche et l’Innovation” under number 2013-0047. EP is authorized to perform experiments on vertebrate animals (license 75–1265) issued by the ‘Direction Départementale de la Protection des Populations de Paris’ and is responsible for all the experiments conducted personally or under his supervision as governed by the laws and regulations relating to the protection of animals.

### Bone marrow-derived macrophages

Bone marrow cell suspensions were recovered from tibias and femurs of female C57Bl/6JRj mice in Dulbecco’s phosphate buffered solution (PBS), and cultured in complete medium containing 4 g/L glucose, 1 mM pyruvate and 3.97 mM L-Alanyl-L-Glutamine and supplemented with 10% heat-inactivated fetal calf serum (FCS), streptomycin (50 µg/mL), and penicillin (50 IU/mL) (Sigma-Aldrich, P4333) and 50 μM β-mercaptoethanol (Sigma-Aldrich, M7522), (de La Llave et al., 2011). Briefly, one million cells per ml of complete medium supplemented with 50 ng/ml of mouse recombinant colony-stimulating factor 1 (rm-CSF1, ImmunoTools) were incubated in bacteriological Petri dishes (Greiner bio-one 664161) at 37°C in 7.5% CO_2_ atmosphere. After 6 days of culture, the medium was removed and adherent cells were incubated with pre-warmed PBS pH 7.4 containing 25 mM EDTA for 30 min at 37°C. Bone marrow-derived macrophages (BMDMs) were detached by gentle flushing, collected, and resuspended in a complete medium supplemented with 30 ng/ml of rmCSF-1. BMDMs were seeded in 100 mm tissue culture dishes (Petri BD Falcon, 353003) for nuclear or total protein extraction and RNA isolation.

### Parasite isolation and macrophage infection

mCherry-transgenic, tissue-derived amastigotes of *Leishmania amazonensis* strain LV79 (WHO reference number MPRO/BR/72/M1841) were isolated from infected footpads of Swiss nude mice (Lecoeur et al., 2020a). After seeding at day 6 of differentiation, BMDMs were left for 5 hours at 37°C to attach to their new culture support. BMDMs were infected with lesion-derived *L. amazonensis* amastigotes at a multiplicity of infection (MOI) of 4 and then cultured at 34°C. These cells were referred to as Infected Macrophages (IMs) versus un-infected BMDMs (UIMs).

### RNA extraction

Large and small (including miRNAs) RNA fractions were isolated from the same macrophage samples using the NucleoSpin miRNA kit (Macherey-Nagel) according to the manufacturer’s instructions. Evaluation of RNA quality was carried out by optical density measurement using a Nanodrop device (Kisker, http://www.kisker-biotech.com). RNAs were isolated from 3 independent biological replicates of UIMs and IMs at 48 hours post-infection (PI).

### RNA-Seq analysis

Library preparation and sequencing was performed at the Biomics platform of Institut Pasteur. mRNA processing: Briefly, DNAse-treated RNA extracts from Large RNA fractions were processed for library preparation using the Truseq Stranded mRNA sample preparation kit (Illumina, San Diego, California) according to the manufacturer’s instructions. An initial poly A+ RNA isolation step (included in the Illumina protocol) was performed with 1 µg of large RNA to isolate the mRNA fraction and remove ribosomal RNA. The mRNA-enriched fraction was fragmented by divalent ions at high temperatures. The fragmented samples were randomly primed for reverse transcription followed by second-strand synthesis to create double-stranded cDNA fragments. No end repair step was necessary. An adenine was added to the 3’-end and specific Illumina adapters were ligated. Ligation products were submitted to PCR amplification. The quality of the obtained libraries was controlled using a Bioanalyzer DNA1000 Chips (Agilent, # 5067-1504) and quantified by spectrofluorimetric analysis (Quant-iT™ High-Sensitivity DNA Assay Kit, #Q33120, Invitrogen). Sequencing was performed on the Illumina Hiseq2500 platform to generate single-end 65 bp reads bearing strand specificity. Reads were cleaned using cutadapt version 1.11 and only sequences at least 25 nt in length were considered for further analysis. STAR version 2.5.0a (Dobin et al., 2013), with default parameters, was used for alignment on the reference genome (GRCm38 from Ensembl database 94). Genes were counted using featureCounts version 1.4.6-p3 (Liao et al., 2014) from the Subreads package (parameters: -t gene -s 0).

miRNA processing: Sequencing was performed on the Illumina Hiseq2500 platform to generate single-end 50 bp reads with strand specificity. Reads were cleaned using cutadapt version 1.11, and only sequences with a length between 18 and 28 nucleotides were considered for further analysis. UMIs were added to read headers using umi-tools extract version 1.0.0 (Smith et al., 2017). Bowtie version 1.2.2 (Langmead et al., 2009), with default parameters, was used for alignment to the reference genome (GRCm38 from Ensembl database 92). Reads were deduplicated using umi-tools dedup to remove genuine PCR duplicates identified by UMIs. miRNAs were counted using featureCounts version v1.6.1(Liao et al., 2014) from the Subreads package (parameters: t miRNA -g Name -s 1) and the annotation from mirBase.

The data are publicly available at the NCBI’s Gene Expression Omnibus repository (ongoing Superseries submission).

RNA-seq data analyses: Count data were analyzed using R version 3.6.1 (R-Core-Team, 2016) and the Bioconductor package DESeq2 version 1.24.0 (Love et al., 2014). The normalization and dispersion estimation were performed with DESeq2 using the default parameters and statistical tests for differential expression were performed applying the independent filtering algorithm. A generalized linear model was set in order to test for the differential expression between the conditions For each pairwise comparison, raw p-values were adjusted for multiple testing according to the Benjamini and Hochberg (BH) procedure (Benjamini and Hochberg, 1995) and genes with an adjusted p-value lower than 0.05 were considered differentially expressed.

### Construction of putative miRNA/mRNA genetic interaction networks

To identify potential miRNA targets, we adopted an approach integrating computational prediction methods and experimental data, based on simultaneous profiling of miRNA and mRNA expression changes in our experimental system. Potential targets of differentially expressed miRNAs were predicted as follows: For each differentially regulated miRNA, putative target mRNAs were first selected based on their inverse, differential expression during infection, considering an adjusted p-value of less than 0.05 as statistically significant for both miRNA and mRNA signals. We next employed the TargetScan (version 7.2) (https://www.targetscan.org/mmu_72/) (Agarwal et al., 2015; McGeary et al., 2019) and miRDB (http://www.mirdb.org/mirdb/index.html) (Chen and Wang, 2020) tools associated with the miRbase database (v22) (https://mirbase.org/search/) (Griffiths-Jones et al., 2006; Kozomara et al., 2019) to screen for interactions between miRNAs and target mRNAs based on sequence complementarity. These putative associations were then visualized utilizing the Cytoscape software package (version 3.10.0) (Shannon et al., 2003), with each node of the predicted miRNA-mRNA networks colored based on the differential expression value (blue, decreased abundance; red, increased abundance).

Secondary structures and thermodynamic stability of prioritized miRNA sequences were predicted using the RNAfold web server (http://rna.tbi.univie.ac.at/cgi-bin/RNAWebSuite/RNAfold.cgi) (Lorenz et al., 2011). Additionally, we employed the RNAcomposer web server (http://rnacomposer.ibch.poznan.pl/) (Popenda et al., 2012) to predict the 3D structure of prioritized miRNAs. Visualization of these three-dimensional structures was performed using the BIOVIA Discovery Studio Visualizer software (https://discover.3ds.com/discovery-studio-visualizer-download). To determine the minimal free hybridization energy of these miRNAs with select potential targets, we utilized the RNAhybrid web server (https://bibiserv.cebitec.uni-bielefeld.de/rnahybrid) (Kruger and Rehmsmeier, 2006).

### Gene ontology and pathway analyses

Gene Ontology (GO) term enrichment analyses were conducted for all mRNAs showing differential expression between uninfected (UIM) and infected (IM) macrophages (p < 0.05), utilizing the GoTermFinder platform (http://www.go.princeton.edu/cgi-bin/GOTermFinder). Subsequently, the REVIGO Web server (http://www.revigo.irb.hr/) was employed to generate GO scatter plots, using the default list size (Medium: 0.7) (Supek et al., 2011). The associated values for each GO term represent the P-value, which is also reflected by the intensity of the bubble color. Bubble size indicates the frequency of the GO term in the underlying GO annotation (GOA) database (bubbles of more general terms are larger). Obsolete GO terms were removed, and SimRel (default) settings was applied to measure semantic similarity.

GO terms enrichment analyses of differentially expressed miRNAs between uninfected (UI) and infected (I) macrophages (p < 0.05) was performed using the miEAA platform (https://ccb-compute2.cs.uni-saarland.de/mieaa/) that integrates data from different databases, such as miRBase, miRWalk et miRTarBase (Backes et al., 2016). miEAA is tailored for miRNA precursors and mature miRNAs from several frequently studied species, such as human, mouse, or rat. Subsequently, GO scatter plots were visualized using the REVIGO Web server as detailed above.

Gene lists used in our analyses were downloaded from the databases KEGG (Kyoto Encyclopedia of Genes and Genomes, https://www.kegg.jp/kegg/) and MGI (Mouse Genome Informatics, https://www.informatics.jax.org/), while the list of experimentally validated transcription factors was used as published by Lambert and colleagues (Lambert et al., 2018). The resulting gene lists were then analyzed for possible miRNAs target sites using the TargetScan platform (version 7.2) (https://www.targetscan.org/mmu_72/), which predicts potential mRNA targets of canonical miRNAs by searching for the presence of 8mer, 7mer, and 6mer sites complementary to the seed region for a given miRNA (Agarwal et al., 2015; Riolo et al., 2020). The 6mer corresponds to a perfect sequence of 6 nucleotides with the seed region of the miRNA (miRNA nucleotides 2 to 7) (Lewis et al., 2005). The most optimal 7mer site, referred to as 7mer-m8, features a match with the seed region strengthened by a pairing with the 8th nucleotide of the miRNA (Brennecke et al., 2005; Krek et al., 2005; Lewis et al., 2005; Lewis et al., 2003). Another 7mer site, 7mer-A1, is also effective as it includes a seed match with an ‘A’ at target position 1 (Lewis et al., 2005). Lastly, the 8mer site consists of a seed match flanked by a match at position 8 and a match with an ‘A’ at position 1 (Lewis et al., 2005). Our miRNA-mRNA interaction networks were further refined using the TargetScan parameter ‘site types’ (8mer, 7mer-m8, 7mer-1A, and non-canonical pairing) and the number of sites in all annotated 3′ UTRs. This parameter selection is based on the notion that among the presented motifs, the greater the number of nucleotides involved in pairing, the more effective the interaction will be. Thus, the type of site (e.g., whether the site is an 8mer or 7mer-m8) strongly influences repression efficacy. The number of sites also influences efficacy, with each additional site typically acting independently to confer additional repression, although sites situated between 8 and 40 nt from each other tend to act cooperatively (Agarwal et al., 2015; Grimson et al., 2007). This approach facilitates direct comparison with other studies employing TargetScan and ensures the reproducibility of our results.

### Ago-APP

Ago-APP (Ago <u>A</u>ffinity <u>P</u>urification by <u>P</u>eptides) was essentially carried out as described earlier in (Hauptmann et al., 2015). Briefly, per 50 μL of anti-FLAG-M2 agarose (Sigma-Aldrich), a minimum of 100 μg FLAG-tagged peptide (TNRC6B 599–683) was coupled to the beads for 3 h at 4 °C. Excess peptide was removed by washing with PBS three times, and lysate was added to the peptide-coupled beads. After incubation for 3 h at 4 °C, the beads were washed four times with NET buffer (50 mM Tris, pH 7.5, 150 mM NaCl, 5 mM EDTA, 0.5 % Nonidet P-40, 10 % glycerol, 1 mM NaF; supplemented with 0.5 mM DTT and 1 mM AEBSF before use) and once with PBS. The beads were eluted by addition of 2× Laemmli sample buffer and incubation at 95 °C for 5 min.

### Phosphosite measurements by mass spectrometry

Phosphosite measurements of Ago2 proteins by mass spectrometry were performed as described earlier in (Quevillon Huberdeau et al., 2017). SDS-gel bands of affinity-purified Ago proteins were excised, transferred into 2ml micro tubes (Eppendorf), washed with 50 mM NH_4_HCO_3_, 50 mM NH_4_HCO_3_/acetonitrile (3/1), 50 mM NH_4_HCO_3_/acetonitrile (1/1) and lyophilized. After reduction and alkylation of cysteines with DTT and Iodoacetamide, gel slices were again washed as described above and lyophilized. Proteins were subjected to *in gel* tryptic digest overnight at 37 °C with approximately 2 μg trypsin per 100 μl gel volume (Trypsin Gold, mass spectrometry grade, Promega). Peptides were first extracted with 100 mM NH_4_HCO3, followed by 50 mM NH_4_HCO_3_ in 50 % acetonitrile. Combined eluates were lyophilized and reconstituted in 20 μl 1 % TFA prior to LC-MS/MS. Separation of peptides by reversed-phase chromatography was carried out on an UltiMate 3000 RSLCnano System (Thermo Fisher Scientific, Dreieich) which was equipped with a C18 Acclaim Pepmap100 preconcentration column (100 μm i.D.x20 mm, Thermo Fisher Scientific) followed by an Acclaim Pepmap100 C18 nano column (75 μm i.d.x250 mm, Thermo Fisher Scientific). A linear gradient of 4 % to 40 % acetonitrile in 0.1 % formic acid over 60 min was used to separate peptides at a flow rate of 300 nl/min. Via a CaptiveSpray nanoflow electrospray source the LC was coupled to a maXis plus UHR-QTOF System (Bruker Daltonics, Bremen). Data-dependent acquisition of MS/MS spectra by CID fragmentation was performed at a resolution of minimum 60000 for MS and MS/MS scans. The MS spectra rate of the precursor scan was 2 Hz processing a mass range between m/z 175 and m/z 2000. Via the Compass 1.7 acquisition and processing software (Bruker Daltonics) a dynamic method with a fixed cycle time of 3 s and an m/z dependent collision energy adjustment between 34 and 55 eV was applied. Raw data processing was performed in Data Analysis 4.2 (Bruker Daltonics). ProteinScape 3.1.3 (Bruker Daltonics) in connection with Mascot 2.5.1 (Matrix Science) facilitated database searching. The following search parameters were applied: database Swiss-Prot *Mus musculus*, enzyme specificity trypsin with 2 missed cleavages allowed, precursor tolerance 0.02 Da, MS/MS tolerance 0.04 Da, variable modifications: acetylation at protein N-terminus, carbamidomethylation and propionamide modification of cysteine, oxidation of methionine, deamidation of asparagine and glutamine, phosphorylation of serine, threonine and tyrosine. Mascot peptide ion-score cut-off was set 15. Phosphopeptide fragment spectra were evaluated manually. Calculation of relative abundances of phosphopeptide species at the S825-S835 phospho-cluster was based on peptide intensities. The sum of peptide intensities of all peptide species detected at the cluster position was considered as 100 %. Peptide intensities of individual phosphorylated or non-phosphorylated species were then used to determine the relative abundance of each peptide as a percentage.

## RESULTS AND DISCUSSION

### *L. amazonensis* infection modulates expression of RISC components and Ago2 phosphorylation

Argonaute (Ago) proteins are fundamental regulators of mammalian gene expression (Mauro et al., 2023). As key components of the RNA-induced silencing complex (RISC), they are essential in mediating the functions of microRNAs (miRNAs), a class of non-coding RNAs that regulate gene expression at the post-transcriptional level promoting mRNA degradation or inhibiting protein translation (Larivera et al., 2023) (Figure 1A). The dysregulation of RISC functions has been implicated in a wide range of pathologies – including microbial infection - revealing miRNAs as promising therapeutic targets for RNA-based intervention strategies (Li et al., 2014). However, how pathogenic microbes exploit the Ago/RISC/miRNA axis to reprogram their host cells is only poorly understood. Here we use the protozoan parasite *Leishmania* as a unique model system to address this major scientific gap and gain mechanistic insight into how these human pathogens manipulate miRNA networks to subvert host defenses and ensure their own survival.

**FIGURE 1:**
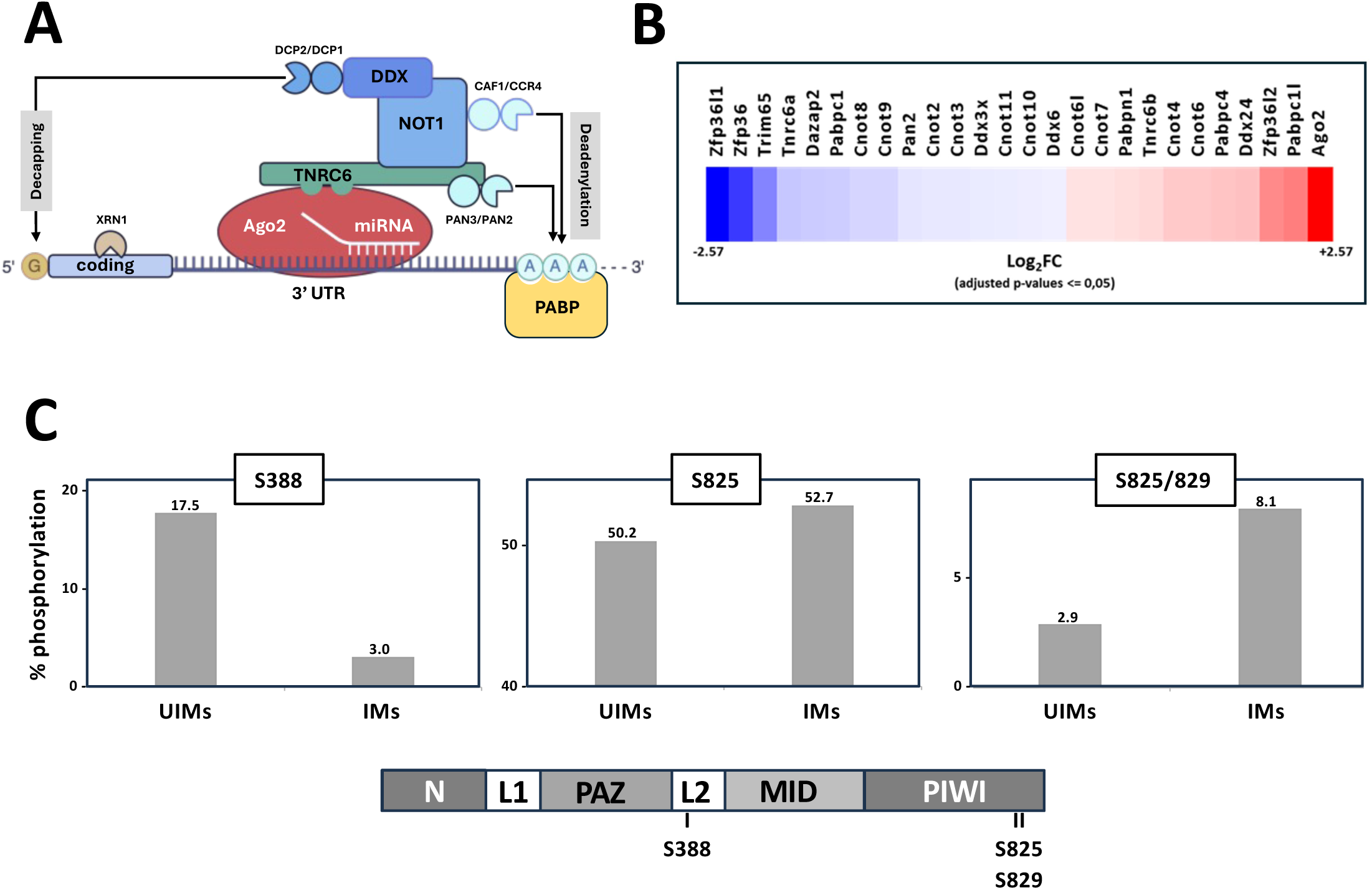
(A) Schematic overview of the Ago2/RISC/miRNA machinery. miRNA-loaded Ago2 (RISC) binds TNRC6 (aka GW182) via the interaction between tryptophan (W) binding pockets in AGO and glycine-tryptophan (GW) repeats in TNRC6. TNRC6 destabilizes target mRNAs by recruiting two deadenylation complexes composed of either CAF1-CCR4-NOT1 or PAN2-PAN3, and by recruiting decapping factors, including the decapping enzyme DCP2. Finally, XRN1 degrades the decapped mRNA in a 5′ to 3′ direction. **(B)** Heat map representing the Log2 Fold Change values between *L. amazonensis*-infected and uninfected BMDMs at day 3 post-infection. Key components of the Ago2/RISC/miRNA machinery are indicated. Note that Ago2 shows a 6-fold induction during infection. **(C)** Phosphorylation analysis of Ago2 protein purified by Ago-APP from *L. amazonensi*s-infected (IMs) vs. uninfected BMDMs (UIMs). LC-MS/MS measurements detected Ago2 phosphorylation changes of serine 388 (left), serine 825 (middle), and serine 825/829 (right). The lower panel shows a schematic view of the Ago2 domain structure, with phosphorylation sites indicated.

Drawing from previously published RNAseq analyses of *L. amazonensis*-infected, bone marrow-derived macrophages (BMDMs), we first investigated expression changes of the RISC members and associated proteins indicated in Figure 1A. Significant expression changes compared to uninfected BMDMs (p < 0.05) were observed for major RISC components (Figure 1B), including (i) the Argonaute family member Ago2 (over 5-fold increase), a multifunctional scaffolding protein that confers miRNA silencing, or (ii) the RNA-binding protein ZFP36L1 binds adenylate-uridylate-rich elements (AREs) in the 3’UTRs of mRNAs, recruiting the CCR4-NOT deadenylase complex to remove poly(A) tails, leading to mRNA degradation (Loh et al., 2020).

Ago proteins are phosphorylated at different residues and specific functions have been assigned to these modifications (Golden et al., 2017; Hauptmann et al., 2015; Quevillon Huberdeau et al., 2017). Whether *Leishmania* infection alters such phosphorylation patterns, however, has not been investigated. Therefore, we isolated all four murine Ago proteins from *Leishmania*-infected BMDMs using the AgoAPP approach (Hauptmann et al., 2015) and analyzed phosphorylation patterns by mass spectrometry (Figure 1C). Indeed, phosphorylation of serine 388 (S388), which was associated with specific sub-cellular Ago localization patterns (Zeng et al., 2008), was strongly reduced upon infection (Figure 1C, left panel). Ago proteins undergo a phosphorylation cycle involving a hypo- and hyper-phosphorylation of a C-terminal Ser/Thr cluster (Golden et al., 2017; Quevillon Huberdeau et al., 2017). Depending on the amount of the negative charge that is added by phosphorylation to this cluster, mRNA binding of Ago/miRNA complex is inhibited. While single phosphorylation events at Ser825 are not significantly changed upon infection (Figure 1C, middle panel), peptides that carry double phosphorylations at S825 and S829 are clearly increased during *Leishmania* infection (Figure 1C, right panel).

In conclusion, *Leishmania* has a profound effect on the expression of RISC components in infected macrophages that on one hand triggers increased expression of Ago2, while on the other hand balancing its functions by Ago2 hyper-phosphorylation to reduced Ago/miRNA activity. These results primed us to directly investigate changes of the miRNA landscape during infection.

### *L. amazonensis* subverts both the mRNA and miRNA landscapes in infected BMDMs

Changes in miRNA and mRNA abundances were investigated in uninfected, bone marrow-derived macrophages (BMDMs, referred to as UIMs) and *L. amazonensis* infected BMDMs (referred to as IMs) established using virulent, lesion-derived amastigotes (Figure 2). Large (mRNAs) and small (primarily miRNAs) RNA fractions were isolated from these samples at day 2 post-infection (PI) and subjected to RNAseq analysis (Figure 2A). As judged by the expression changes observed across three independent biological replicates, *Leishmania* infection has a major impact on both the mRNA and miRNA profiles of IMs, with significant changes observed for the abundance of 5742 mRNAs (10.36 % of total, padj <0.05) and 102 miRNAs (6.04 % of total, padj <0.05) (Figure 2B1 and 2C1). Inspection of biological functions enriched in the differentially expressed genes (Figure 2B2) and miRNAs (Figure 2C2), revealed a series of GO terms that were previously associated with *Leishmania*-mediated immune subversion and infected macrophages and that converged in both the miRNA and mRNA datasets, including GO terms linked to inflammatory response, NF-κB signaling, apoptosis, or essential metabolic processes related to cholesterol and fatty acid biosynthesis (Figures 2B2 and C2).

**FIGURE 2:**
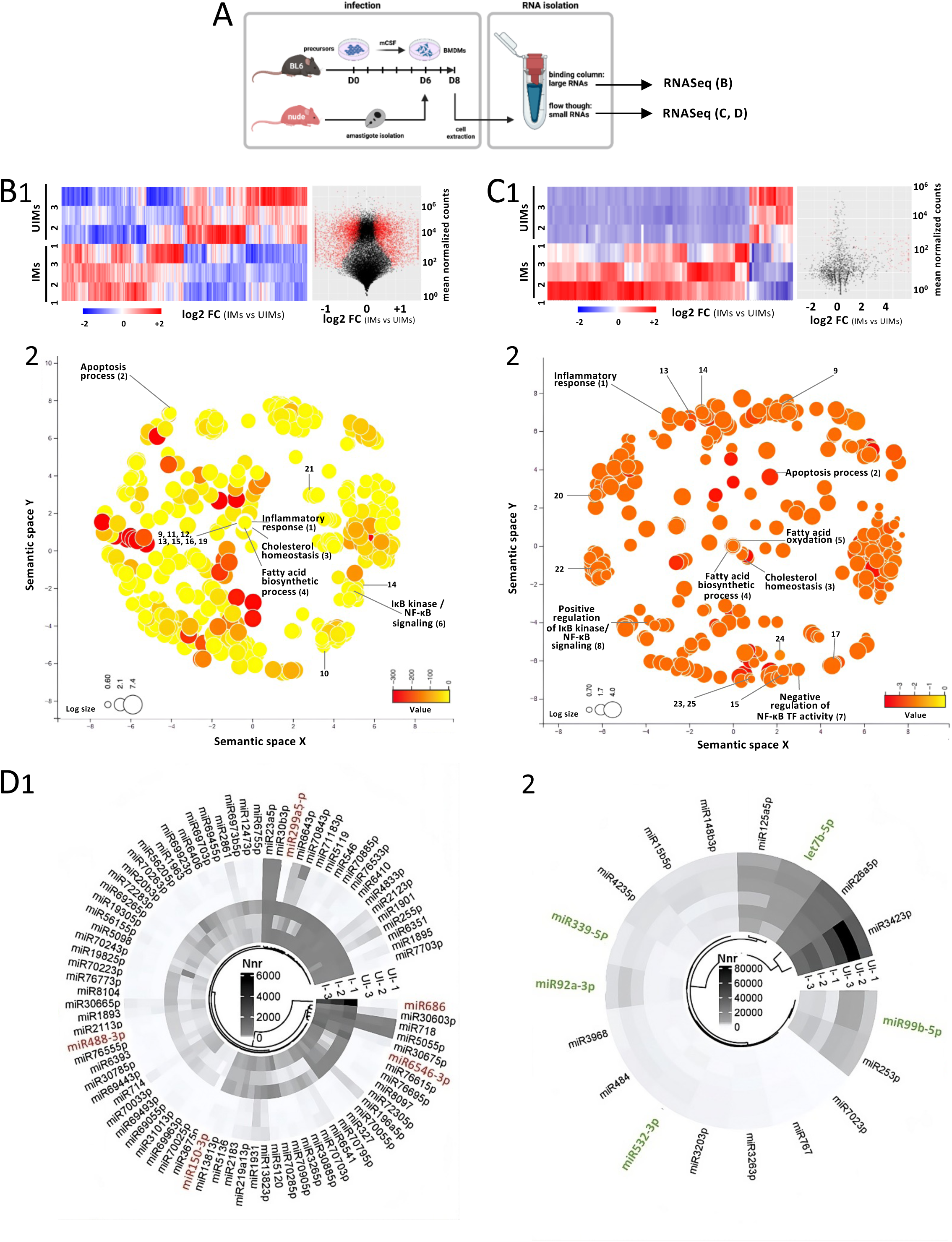
Dual RNAseq analysis of mRNA and miRNA fractions in *L. amazonensis-*infected versus non-infected macrophages. **(A)** Experimental pipeline for the generation of BMDMs, their infection, and isolation of large versus small RNAs. BMDMs were derived from mCSF-treated C57BL/6 bone marrow progenitors and were infected (IMs) or not (UIMs) with lesion-derived amastigotes of *L. amazonensis* for 2 days. Large and small RNAs were isolated from the same samples (n=3 independent experiments) and subjected to RNAseq analysis. **(B, C)** Heat maps, volcano plots and REVIGO enrichment maps showing respectively differentially expressed mRNAs and miRNAs across three biological replicates (1-3) in uninfected (UIMs) and infected (IMs) macrophages (B1 and B2, left panels, log2 fold changes (FC) are shown), the distribution of differentially expressed mRNAs and miRNAs according to their average, normalized read counts (B1 and C1, right panels, statistically significant signals with p < 0.05 are indicated in red), and biological functions enriched in the datasets of differentially expressed mRNA (B2) and miRNA (C2). Bubble color reflects the p-value of the enrichment and bubble size indicates the frequency of the GO term in the underlying GOA database. For space reasons, only few bubbles are labeled with their corresponding GO term, while others are numbered as follows: 1, Inflammatory response (GO:0006954); 2, Apoptosis process (GO:0006915); 3, Cholesterol homeostasis (GO:0042632); 4, Fatty acid biosynthetic process (GO: 0006633); 5, Fatty acid oxidation (GO: 0019395); 6, I-κB kinase/ NF-κB signaling (GO:0007249); 7, Negative regulation of NF-κB TF activity (GO:0032088); 8, Positive regulation of I-κB kinase/ NF-κB signaling (GO:0043123); 9, Response to LPS (GO: 00032496); 10, Response to protozoan (GO:0001562); 11, Macrophage activation (GO: 0042116); 12, Cytokine production involved in immune response (GO:0002367); 13, JNK cascade (GO:0007254); 14, MAPK cascade (GO:0000165); 15, Regulation of gene expression (GO: 0010468); 16, Regulation of DNA-binding TF activity (GO:0051090); 17, Positive regulation of DNA binding (GO:0043388); 19, Chromatin remodelling (GO:0006338); 20, Histone H4 acetylation (GO:0043967); 21, Epigenetic regulation of gene expression (GO:0040029); 22, mRNA transport (GO:0051028); 23, RNA-mediated gene silencing (GO:0031047); 24, Positive regulation of miRNA-mediated gene silencing (GO:0043388); 25, miRNA-mediated gene silencing by inhibition of translation (GO:0035278). **(D)** Circular heatmaps of miRNAs showing increased (D1) or decreased (D2) normalized read counts during infection. Highlighted in red and green are miRNAs with high similarity to human miRNAs (except miR-686 and miR6546-3p). Nnr, Normalized read counts.

Closer inspection of miRNA read depth variations during infection identified 84 miRNAs (82.4%) with and 18 miRNAs (17.6%) with decreased abundance, suggesting that intracellular *L. am* has a profound effect on the miRNA landscape in agreement with a previously published report using *L. amazonensis* infected BMDMs from BALB/c (Muxel et al., 2017) (Figure 2D1 and D2, respectively). Important read depth variations were observed for mmu-miR-686 that showed an over 1000-fold increase in abundance (3 normalized reads in UIMs vs 3953 in IMs), as well as for mmu-miR-6546-3p (2 vs 891, 445-fold increase) and mmu-miR-718 (261vs 1,023, 3.9-fold increase) (Figure 2D1). Significant increase in expression was further observed for a series of murine miRNAs with high homology to human (https://rnacentral.org/), such as mmu-miR-150-3p, mmu-miR-488-3p, and mmu-miR-299a-5p. In contrast, the strongest reduction in read depth during infection was observed for mmu-miR-342-3p (55,522 vs 31,602), mmu-let-7b-5p (11,428 vs 7,176), mmu-miR-99b-5p (4,508 vs 1,926), mmu-miR-92a-3p (2,278 vs 611), mmu-miR-339-5p (1,271 vs 469), and mmu-miR-532-3p (741 vs 394). Decreased abundance of mmu-let-7b-5p has been also observed during infection with *L. infantum* and *L. major* in human macrophages (Geraci et al., 2015; Ramos-Sanchez et al., 2022), suggesting a key role of this miRNA in intracellular parasite infection.

In conclusion, our data confirm the differential expression of miRNAs during intracellular *Leishmania* infection, which correlated with important expression changes at mRNA levels, suggesting the existence of regulatory miRNA-mRNA networks that may govern the establishment of an immune-compromised host cell phenotype. We next applied a series of computational tools to further assess possible miRNA-mRNA regulatory interactions.

### Modelling interaction networks related to reduced miRNA expression

miRNA-mRNA regulatory networks are characterized by reciprocal expression changes between both types of RNA, where increased miRNA usually causes decreased expression of its target mRNA (and *vice versa*). We investigated the existence of such networks by first focusing on down-regulated miRNAs whose expression changes are supported by at least 50 normalized reads, and their predicted mRNA targets that show increased abundance during infection (Log2 Fold Change (FC) ≥ 1, p < 0.05). Only 2 out of the 18 downregulated miRNAs did not pass this filtering step and were excluded (i.e. mmu-702-3p, mmu-miR-99b-5p). Down-regulation of the remaining 16 miRNAs were tightly associated with increased abundance of their predicted target mRNAs as illustrated by the interaction network shown in Figure 3A. Important differences were observed between these networks both with respect to the level of miRNA down-regulation as well as the number of possible target mRNAs (Figure 3A). For example, mmu-miR-532-3p and mmu-miR-342-3p respectively are predicted to target only 5 and 10 transcripts, while on the other extreme, mmu-484 and mmu-767 respectively are predicted to target 136 and 97 transcripts (Figure 3A). Some sets of upregulated transcripts are predicted to be solely under the control of a single miRNA, as illustrated by mmu-767 (74 transcripts), mmu-484 (36 transcripts), mmu-miR-3968 (22 transcripts), and mmu-let-7b-5p (8 transcripts). Thus, affecting a single miRNA during intracellular *Leishmania* infection can generate a pleiotropic outcome with important consequences on cellular processes and diverse pathologies (Valinezhad Orang et al., 2014).

**FIGURE 3:**
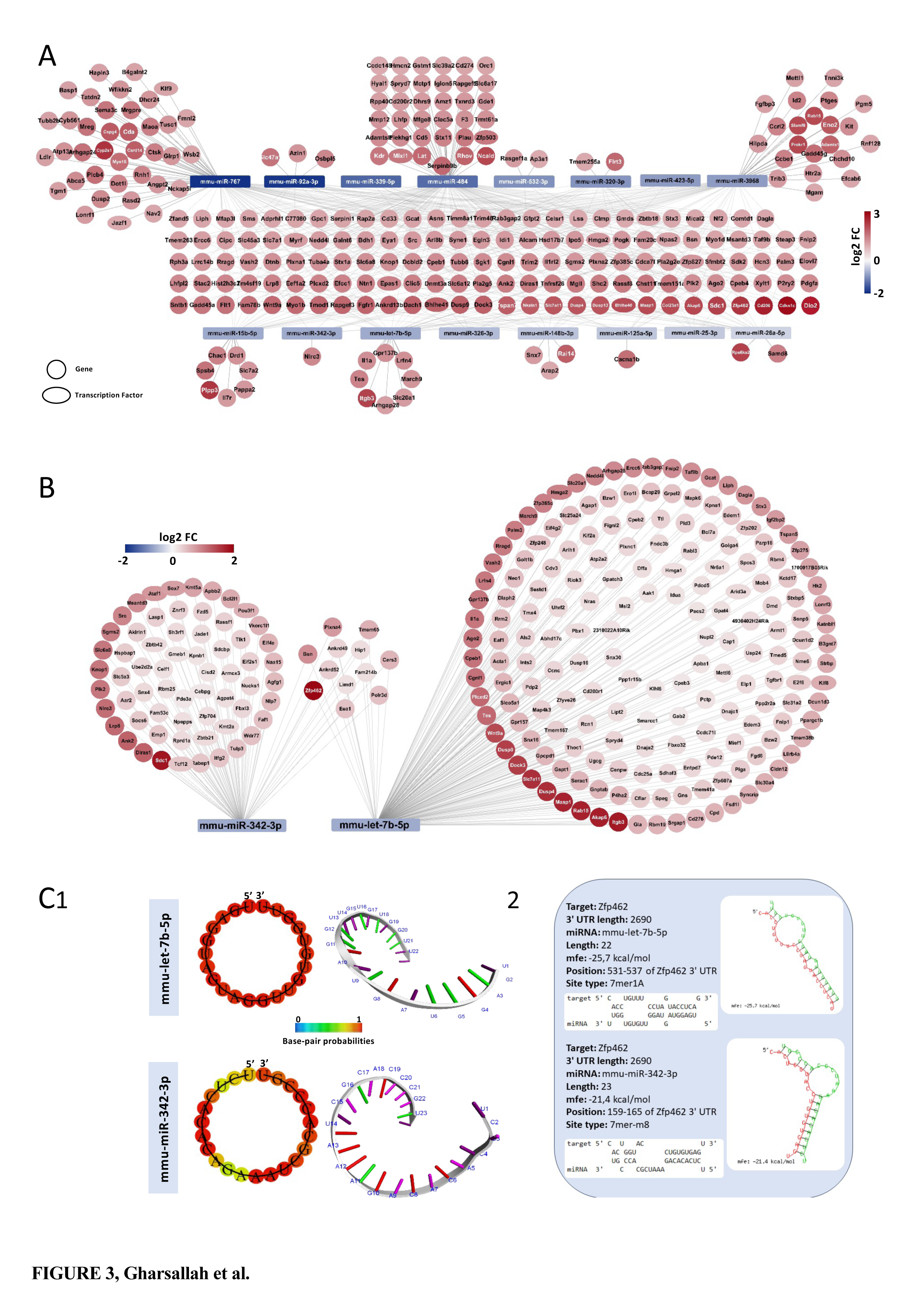
Gene regulatory networks between down-regulated miRNAs and putative target mRNAs showing increased abundance in infected macrophages. **(A)** Predicted interactions are shown between miRNAs (rectangles, fill color indicates level of decreased abundance in log2 FC) and mRNAs (circles, fill color indicates level of increased abundance in log2 FC). Transcription factors are indicated by the oval shape. Shown interactions are based on miRNAs showing a difference of 50 or more normalized reads between infected (IMs) and uninfected (UIMs) groups, and mRNAs showing IMs vs UIMs expression changes of Log2 FC ≥ 1. **(B)** Predicted interaction maps between down-regulated mmu-let-7b-5p and mmu-miR-342-3p and their potential target mRNAs showing increased abundance in IMs vs UIMs. **(C)** 2D and 3D structures of mmu-let-7b-5p and mmu-miR-342-3p (C1). The probability of miRNA binding to the putative target mRNA *Zfp462* is indicated by the color. Target position, length, site type and minimum hybridization energy (mfe) are indicated in panel C2.

We next focused on the two miRNAs mmu-let-7b-5p (11,428 in UIMs vs 7,176 normalized read counts in IMs, Log2FC = -0.752) and mmu-miR-342-3p, based on (i) their significant differential read count between UIMs and IMs, (ii) their conservation to human miRNAs (Pasquinelli et al., 2000), https://rnacentral.org/sequence-search/?jobid=c25cc34b-01cf-42ae-a24e-611a1132aa1e), and their involvement in key biological processes, including apoptosis, Jak/Stat signaling, or NF-κB signaling (Cui et al., 2019; Lecoeur et al., 2022b; Taghehchian et al., 2023; Tai et al., 2015; Thammaiah and Jayaram, 2016). Consolidating our target mRNA predictions from TargetScan and miRDB platforms established a list of 192 putative mmu-let-7b-5p mRNA targets showing increased abundance during infection, including *Integrin Beta 3 subunit* (*Itgb3*, Log2 FC = 2.35), *Argonaute RISC Catalytic Component 2* (*Ago2*, Log2 FC = 1.37), *Dual-specificity phosphatase 9* (*Dusp9*, Log2 FC = 1.94) (Figure 3A) and 7 mRNAs encoding for transcription factors (TFs) or epigenetic regulators (ERs), *i.e*. *Arid3a*, *E2f6*, *Hmga1*, *Klf8*, *Nr6a1*, *Pbx1*, and *Zfp202* (Figure 3B), which are involved in regulation of proliferation regulation, cell differentiation, DNA structure modification, and cancer-related signaling pathways (Bai et al., 2024; Guo et al., 2023; Kao et al., 2024; Patterson et al., 2008; Trimarchi et al., 2001; Wang et al., 2022; Zhang et al., 2024).

Likewise, decreased expression of mmu-miR-342-3p was associated with increased expression during infection of 75 putative mRNA targets, including *Syndecan 1* (*Sdc*1, Log2 FC = 2.37), *LDL Receptor Protein 8* (*Lrp*8, Log2 FC = 2.82) (Figure 3A) and 9 TFs/ERs (*Cebpg*, *Gmeb1*, *Jazf1*, *Kmt2a*, *Pou3f1*, *Sox7*, *Tcf12*, *Zbtb21*, and *Zbtb42*) (Figure 3B), which regulate genes involved in several biological processes such as glucocorticoid synthesis, regulated cell death or proteasomal protein degradation (An et al., 2019)

Both mmu-let-7b-5p and mmu-miR-342-3p share 12 predicted targets, including *Zinc Finger Protein 42* (*Zfp42*) (Log2 FC= 2.537), which shows high base pairing probability that suggests strong binding of both miRNAs (Figure 3C1). Specific interaction between both miRNAs and *Zfp42* mRNA is further supported by the low hybridization energy below -20 kcal/mol and the presence of strong 7mer-m8 and 7mer1A binding properties (Figure 3C2).

In conclusion, our analyses reveal the presence of putative miRNA-mRNA networks that may participate in increased gene expression observed during *Leishmania* infection. The identification of *Ago2* as a putative miRNA target further suggests the existence of a regulatory feedback loop during *Leishmania* infection, where miRNAs can affect their own production as seen recently in mammalian development (Liu et al., 2021). Furthermore, the identification of TFs as putative miRNA targets opens the possibility to control macrophage gene expression in a cooperative fashion through feed-forward loops (FFLs) and feedback loops (FBLs) (Shalgi et al., 2007; Tsang et al., 2007).

### Modelling of interaction networks related to increased miRNA expression

We next applied the same filtering procedure as indicated above to establish networks between the 84 miRNA showing increased expression during infection and the 199 down-regulated mRNAs (Log2 FC ≤ -0.5). The 33 miRNAs of the network shown in Figure 4A passed this filtering step and were prioritized for network analysis considering the level of their increased expression and their high conservation with human miRNA for some of them. As expected, increased expression of these miRNAs correlated with decreased abundance of their predicted mRNA targets (Figure 4A). Important differences were observed for the number of putative mRNA targets that are predicted to interact with these miRNAs (Figure 4A), with for example mmu-miR-299a-5p predicted to target only *Ring Finger Protein 150* (*Rnf150*), or mmu-miR-6546-3p to target just 9 transcripts (including 1 TF), while mmu-miR-7661-5p is predicted to target 93 transcripts (including 6 TFs) (Figure 4A).

**FIGURE 4:**
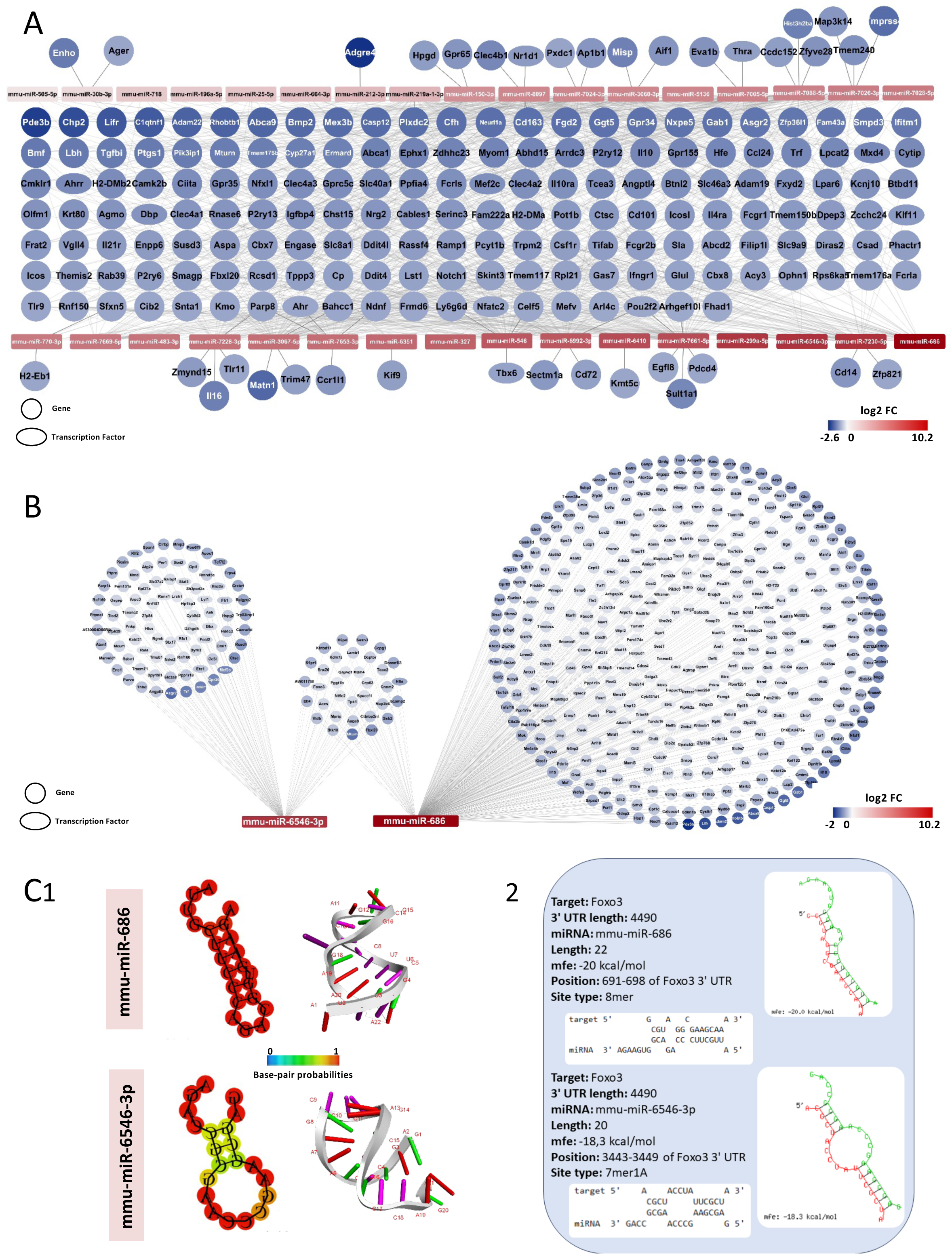
Gene regulatory networks between up-regulated miRNAs and putative target mRNAs showing reduced abundance in infected macrophages. **(A)** Predicted interactions are shown between miRNAs (rectangles, fill color indicates level of increased abundance in log2 FC) and mRNAs (circles, fill color indicates level of decreased abundance in log2 FC). Transcription factors are indicated by the oval shape. Shown interactions are based on miRNAs showing a difference of 50 or more normalized reads between infected (IM) and uninfected (UIMs) groups, and mRNAs showing IMs vs UIMs expression changes of Log2 FC ≤ -1. **(B)** Predicted interaction maps between up-regulated mmu-686 and mmu-miR-6546-3p and their potential target mRNAs showing decreased abundance in IMs vs UIMs. **(C)** 2D and 3D structures of mmu-686 and mmu-miR-6546-3p (C1). The probability of miRNA binding to the putative target mRNA *Foxo3* is indicated by the color. Target position, length, site type and minimum hybridization energy (mfe) are indicated in panel C2.

For subsequent analyses, we focused our attention on two highly expressed miRNAs. First, mmu-miR-686 shows an over 1000-fold increase of abundance during infection and is predicted to target 433 mRNAs that showed statistically significant reduced abundance during infection (p < 0,05). Putative mRNA targets showing reduced abundance in our analysis include the mRNA decay-activating protein *Zinc Finger Protein 36*, C3H Type (*Zfp36l1*, Log2 FC = -3.12), *Ribonuclease A Family Member K6* (*Rnase6*, Log2 FC = -2.4) or *Caspase 12* (*Casp12*, Log2 FC = -3.34) (Figure 4B) linking this miRNA to key biological processes such as apoptosis, cell proliferation and chromatin binding. MiR-686 has also the potential to negatively regulate the transcripts of 31 transcription factors, including *Elf4*, *Bcl6*, *Stat1*, *Foxo3*, *Hmbox1*, and *Bcl6* (Figure 4B), adding cell proliferation, differentiation, apoptosis and antimicrobial resistance to its putative list of target processes (Lecoeur et al., 2022a; Spath et al., 2009; Xu et al., 2024).

Second, mmu-miR-6546-3p shows an over 200-fold increase of abundance during infection and is predicted to target 117 mRNAs that showed statistically significant reduced abundance during infection (p < 0.05) (Figure 4B), including *G-protein-coupled receptor 35* (*Gpr35*, Log2 FC = -2.54), which facilitates the activity of the C-X-C chemokine receptor (Boleij et al., 2021) or the developmental TFs *MADS box transcription enhancement factor 2* (*Mef2c*, Log2 FC = -2.48) and *transformation-related protein 53-induced nuclear protein 1*, (*Trp53inp1*, Log2 FC = -1.92) (Figure 4B). mmu-miR-6546-3p has further the potential to reduce the expression of 14 transcription factors, including *Ets1*, *Foxo3* (implicated in cell death regulation, (Litvak-Greenfeld and Benhar, 2012), *Mef2c*, *Rela*, *Stat2*, and *Stat3* (Figure 4B), some of which were previously linked to *Leishmania* infection, such as *Stat3* (Mishra et al., 2023) or *Rela,* whose mRNA abundance was reduced during *in vitro* and *in vivo L. amazonensis* infection, dampening the TLR / NF-κB / NLRP3 pathway (Lecoeur et al., 2020a).

Both miRNAs have the potential to synergize for the negative regulation of 34 common mRNA targets, including the TFs *Maturin, Neural Progenitor Differentiation Regulator Homolog* (*Mturn*, Log2 FC = -1.559) (Figure 4B). These putative interactions are further supported by (i) strong miRNA-mRNA base pairing probability and an elevated overall thermodynamic free energy of -5.41 (mmu-miR-686) and -1.97 kcal/mol (mmu-miR-6546-3p) (Figure 4C1). In general, miRNAs with high thermodynamic stability are considered crucial for efficient transcriptional regulation, which explains their high intracellular concentrations (>1,000 molecules per cell) according to recent studies (Denzler et al., 2016; Luna et al., 2015). Furthermore, as exemplified by the putative *Foxo3* mRNA target, both miRNAs show a low hybridization energy below -18 kcal/mol and the presence of strong 8mer and 7mer1A binding properties (Figure 4C2).

In conclusion, just as observed for the miRNA^down^/mRNA^up^ regulatory networks, the miRNA^up^/mRNA^down^ regulatory networks described here propose miRNAs as important pleiotropic regulators that may be engaged by *Leishmania* to efficiently reprogram the host cell expression profile in favor of intracellular parasite survival.

### *In silico* assessment of miRNA-mediated subversion of the TF landscape

One striking signal emerging from our network analyses is the number of differentially expressed TFs and epigenetic regulators (ERs) that are predicted targets of miRNAs. A focused *in silico* analysis of these interactions is shown in Figure 5 that visualizes simplified networks between 15 down-regulated miRNAs and 79 upregulated TF/ER mRNAs (Figure 5, panel A), and *vice versa* between 34 upregulated miRNAs and 160 downregulated, putative TF/ER targets (Figure 5, panel B). These interactions may represent a very cost-efficient parasite immune subversion strategy, where changes in a small set of miRNAs have important consequences on macrophage expression profile and phenotype, either by directly affecting TFs/ERs or by exploiting regulatory feed-forward and feedback loops between miRNAs and TFs/ERs that could jointly control the transcriptomic reprogramming we previously described in *L. amazonensis* infected macrophages and dendritic cells (Lecoeur et al., 2022a; Lecoeur et al., 2020a; Lecoeur et al., 2020b).

**FIGURE 5:**
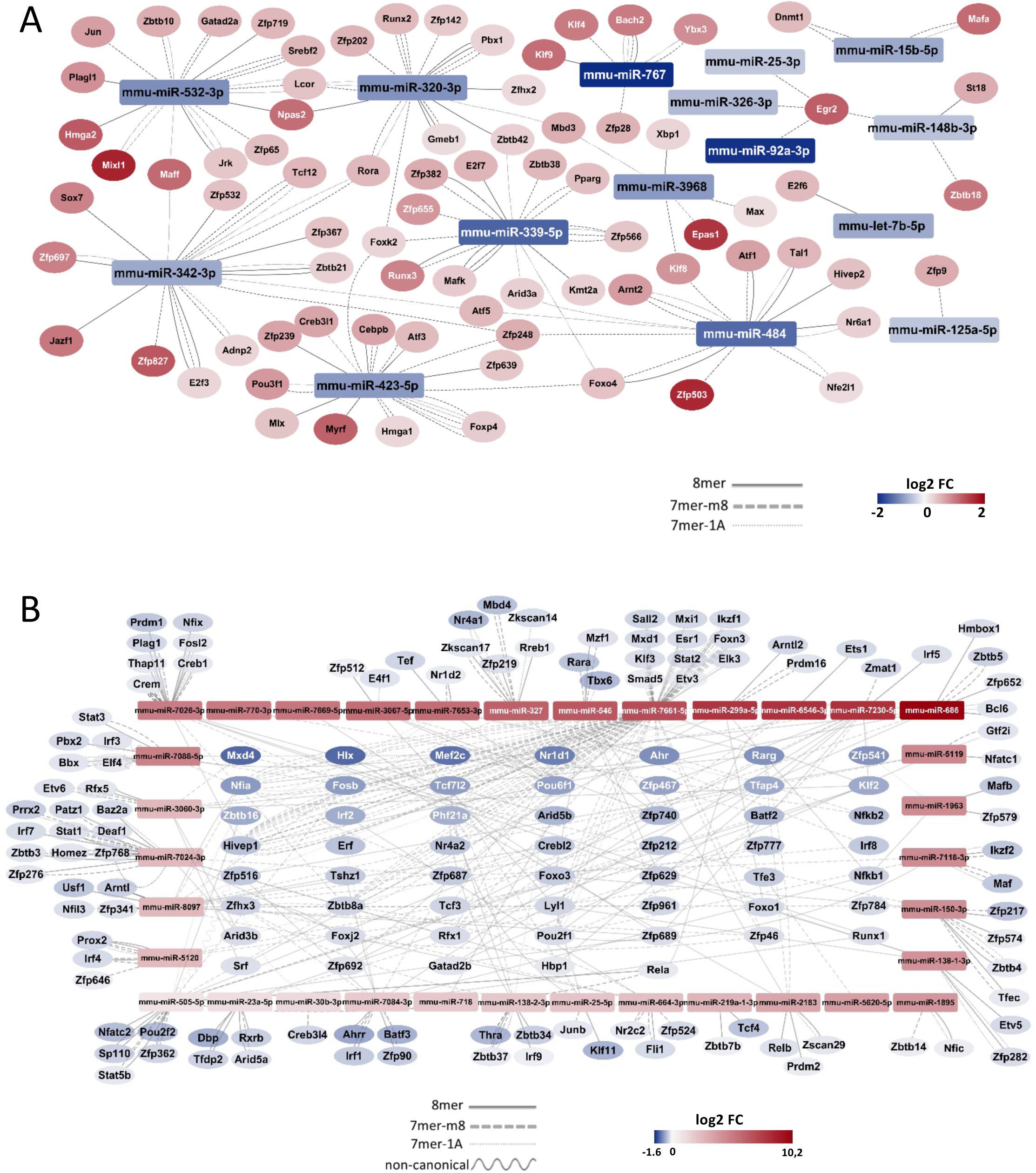
miRNA network analyses focused on putative mRNA targets implicated in expression regulation. mRNAs encoding for transcription factors (TFs) and epigenetic regulators (ERs) showing significantly increased and decreased abundances (padj < 0.05) are shown in red (panel A) and blue (panel B), respectively. The solid lines (8mer) represent the most efficient interaction. Next are the dashed lines (7mer-m8) and dotted lines (7mer-1A), followed by the sine wave lines (non-canonical), indicating varying levels of interaction efficiency.

### *In silico* assessment of miRNA-mediated suppression of the pro-inflammatory response

We have previously identified the TLR / NF-κB axis as a prime target for epigenetic subversion of the macrophage inflammatory response by *L. amazonensis* infection that correlated with histone H3 hypo-acetylation and hypo-methylation (Lecoeur et al., 2020a). Our data here uncover that modulation of the host miRNA landscape may constitute an additional epigenetic layer subverting pro-inflammatory signaling during infection. The focused network shown in Figure 6 identifies 16 down-regulated miRNAs that are predicted to target 16 up-regulated members of this pathway (panel A), while 28 up-regulated miRNAs are predicted to affect 24 down-regulated pathway members (panel B). Even though this one-on-one regulation may indicate only limited influence of miRNA on expression of these targets, placing these interactions into the context of the entire pathway reveals that miRNAs could affect every step of TLR / NF-κB pathway establishing the anti-inflammatory phenotype characteristic of *Leishmania*-infected macrophages (Figure 6C). miRNA^up^/mRNA^down^ interactions are observed for a number of activating pathway members, from surface receptors that trigger the initial activation (e.g. miR-138-2-3p for *Cd40*, miR-7088-5p for *Tnfrsf1a*), down to NF-κB family members (miR-6546-3p for *Rela*, miR-546 for *Relb*, miR-299a-5p for *NFkb1*, miR-327 for *Nfkb2*) that induce inflammatory gene expression. On the contrary, miRNA^down^/mRNA^up^ interactions are observed for deactivating pathway members, such as *Tnfaip3* precited to be targeted by miR-92a-3p. These data reveal miRNAs as potential regulators of the dual inhibition of the macrophage pro-inflammatory response we have described previously (Lecoeur et al., 2020a).

**FIGURE 6:**
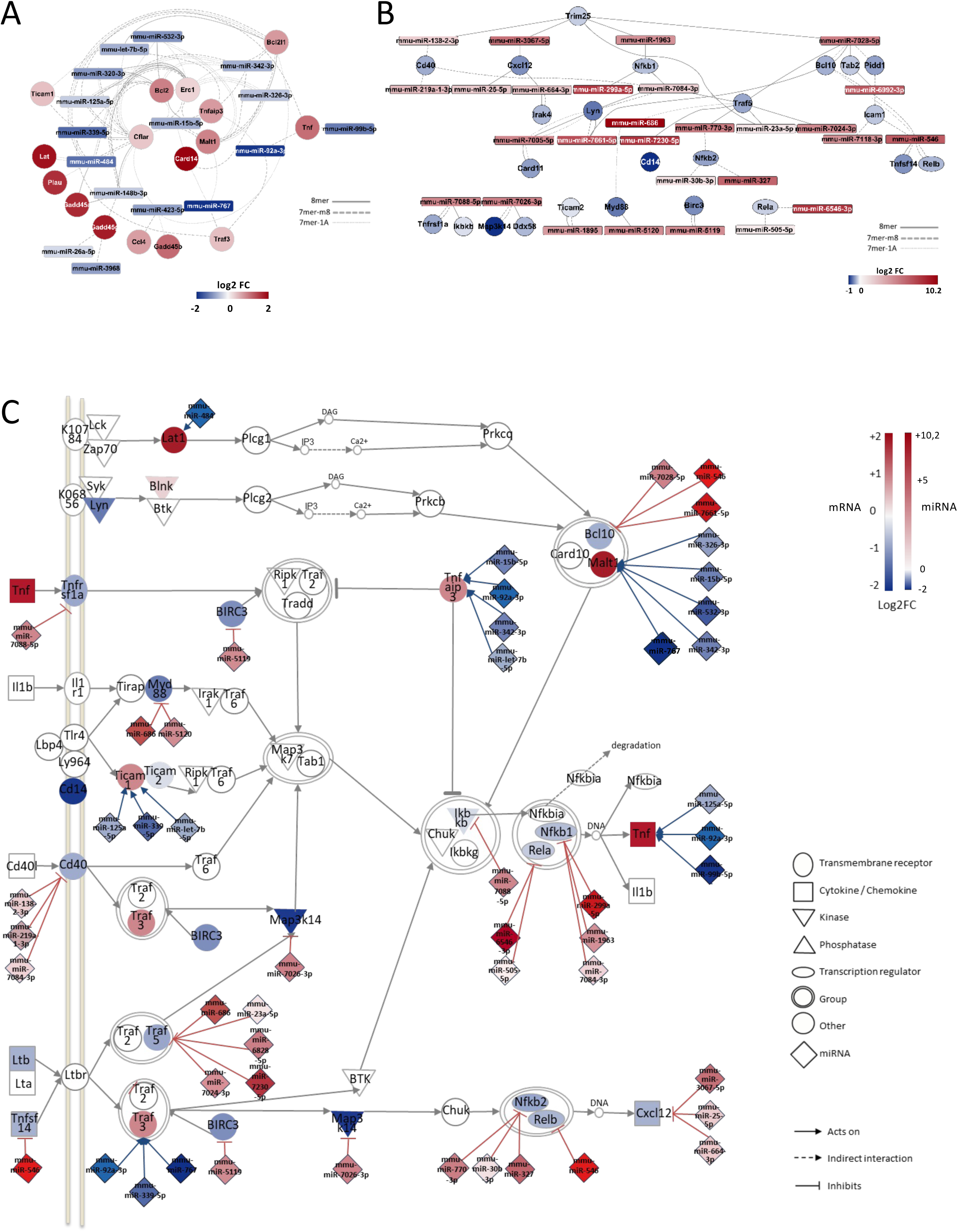
miRNA network analyses focused on putative mRNA targets implicated in the NF-κB pathway. mRNAs encoding for NF-kB pathway members showing significantly increased and decrease (padj < 0.05) mRNA abundances are shown in red (panel A) and blue (panel B), respectively. The solid lines (8mer) represent the most efficient interaction. Next are the dashed lines (7mer-m8) and dotted lines (7mer-1A), indicating varying levels of interaction efficiency. (C) Projection of miRNA and mRNA expression values and putative regulatory interactions on the NF-κB pathway.

### *In silico* assessment of miRNA-mediated induction of cholesterol biosynthesis

Cholesterol biosynthesis represents one of the key biological processes induced during *L. amazonensis* infection believed to be essential for parasite development and vacuole membrane biogenesis (Gutierrez Sanchez et al., 2023; Osorio y Fortea et al.). Our data suggest miRNAs as possible key regulators of this pathway during *Leishmania* infection. The focused network shown in Figure 7 identifies 8 down-regulated miRNAs that are predicted to target 32 up-regulated members of this pathway (panel A), while 20 up-regulated miRNAs are predicted to affect 20 down-regulated pathway members (panel B). Placing these putative interactions in the context of the cholesterol pathway (panel C) sheds a more integrative light on the outcome of these transcriptional modulations and again proposes a dual regulation underlying increased cholesterol production. Certain miRNAs act as possible positive regulators of this pathway, with for example down-regulation of miR-532-3p correlating with upregulation of its mRNA target Sterol Regulatory Element Binding Transcription Factor 2 (*Srebf2*), a master regulator of cholesterol biosynthesis (Amemiya-Kudo et al., 2002). In contrast, miRNA^up^/mRNA^down^ interactions seem to be linked to cholesterol homeostasis, as indicated by down-regulation of the ATP binding cassette subfamily A member *Abca1* (putative target for mmu-miR7661-5p) that functions as a cholesterol efflux pump (Quazi and Molday, 2013), or the cytochrome P450 superfamily member CYP27A1 (putative target for mmu-miR664-3p) controlling cholesterol removal (Pikuleva et al., 1998).Thus the combined effect of these miRNAs likely enhances activating components of cholesterol biosynthesis while at the same time limiting cholesterol removal.

**FIGURE 7:**
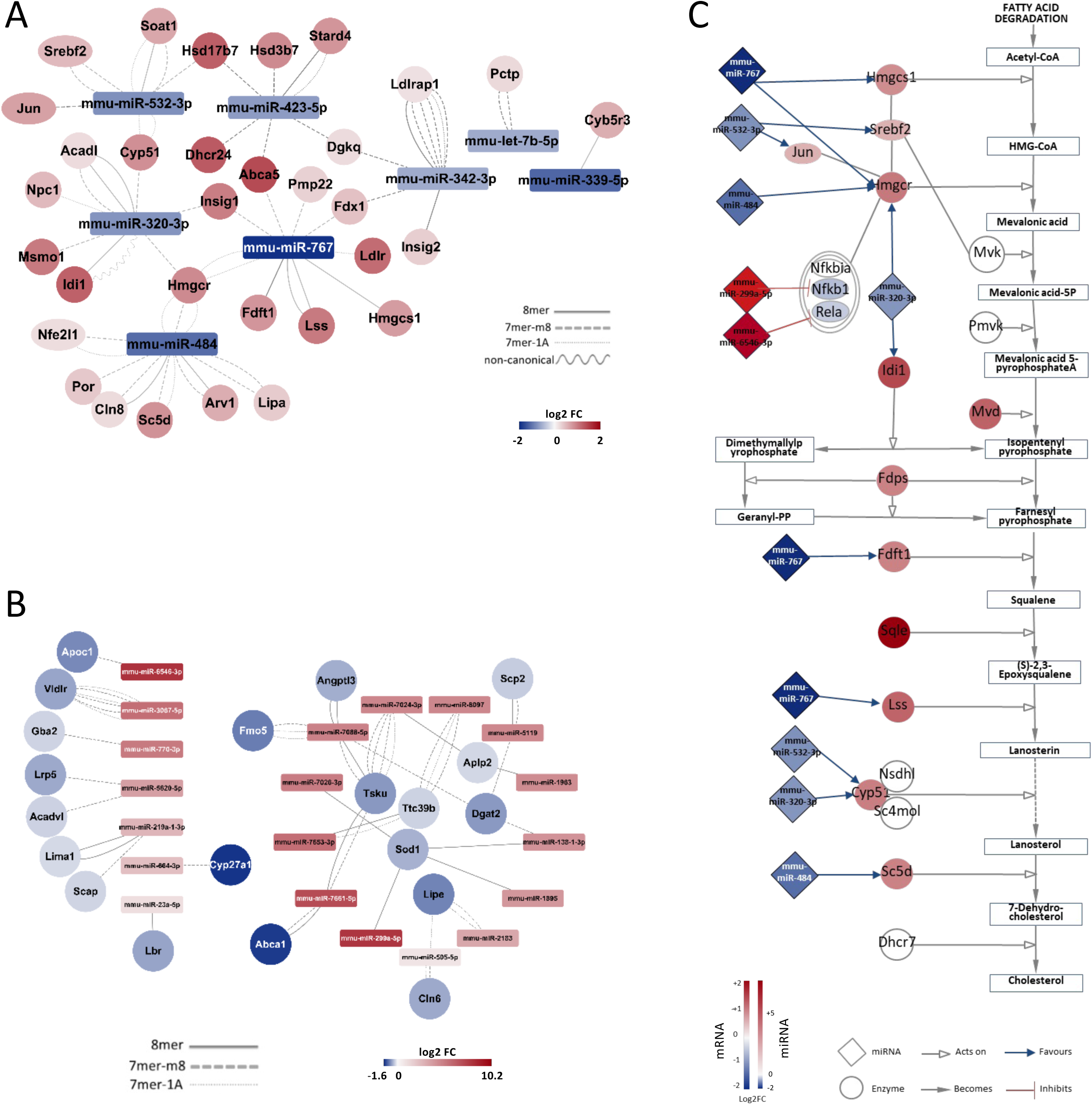
miRNA network analyses focused on putative mRNA targets implicated in cholesterol biosynthesis and homeostasis. mRNAs encoding for proteins implicated in cholesterol production showing significantly increased and decrease (padj < 0.05) mRNA abundances are shown in red (panel A) and blue (panel B), respectively. The solid lines (8mer) represent the most efficient interaction. Next are the dashed lines (7mer-m8) and dotted lines (7mer-1A), indicating varying levels of interaction efficiency. (C) Projection of miRNA and mRNA expression values and putative regulatory interactions on the cholesterol biosynthetic pathway.

## CONCLUSION

Applying dual RNAseq analysis on mRNA and miRNA fractions obtained from *L. amazonensis*-infected and uninfected BMDMs allowed us unprecedented insight into the possible role of the host cell noncoding RNome as a novel target for parasite immune subversion. Based on our computational analyses, the observed miRNA expression changes are predicted to affect the abundance of a large array of transcripts that establish the immune-metabolomic expression profile and cellular phenotype characteristic of *L. amazonensis* infected macrophages (Zhang et al., 2022). The challenge for future studies lies on experimentally validating these interactions for example using RNA pull down experiments to establish defined miRNA-mRNA target relationships or applying genetic ablation on key miRNAs to study their effect on host cell gene expression and intracellular parasite survival. These future investigations will hold great promise to define miRNAs as novel, host-directed therapeutic targets against *Leishmania* using agomir and antagomir strategies to restore the anti-microbial potential of host macrophages (Chong et al., 2021; Murdaca et al., 2019; Rupaimoole and Slack, 2017).

