## Supplementary figures and images for "MAPPING CHANGES OF MIRNA-MRNA NETWORKS IN *LEISHMANIA-INFECTED* MACROPHAGES PREDICTS REGULATORY MIRNA-TF LOOPS AS NOVEL TARGETS OF PARASITE IMMUNE SUBVERSION"

### supplementary figures

Supp Fig

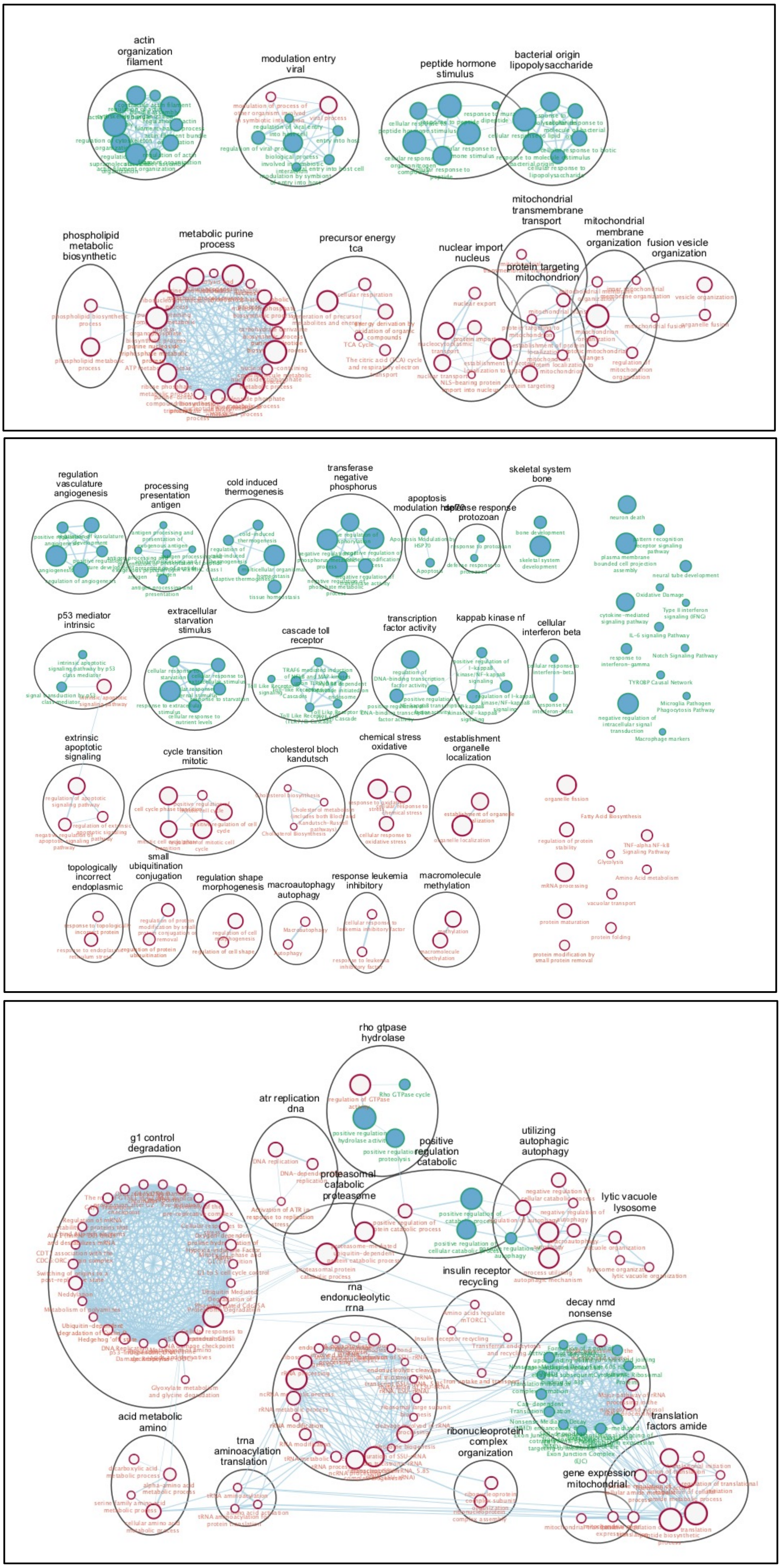

#### mmu-let-7b-5p & mmu-miR-342-3p

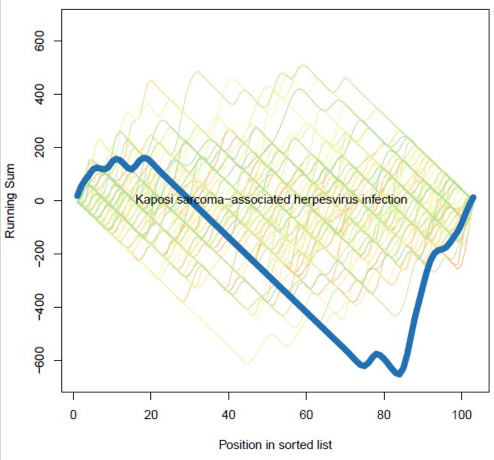


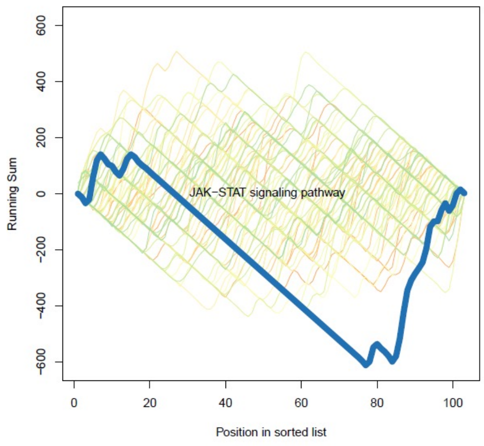


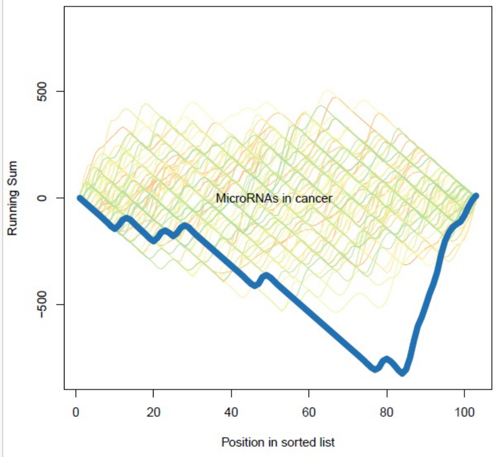


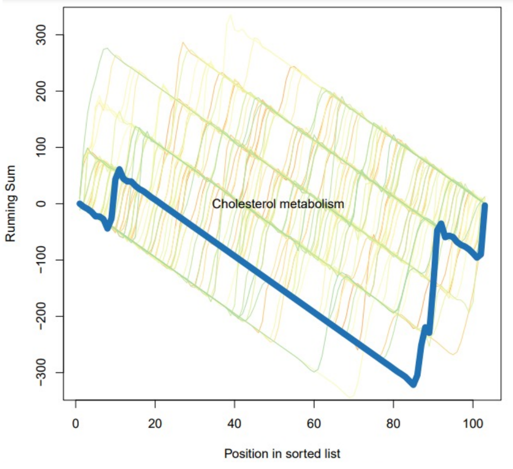


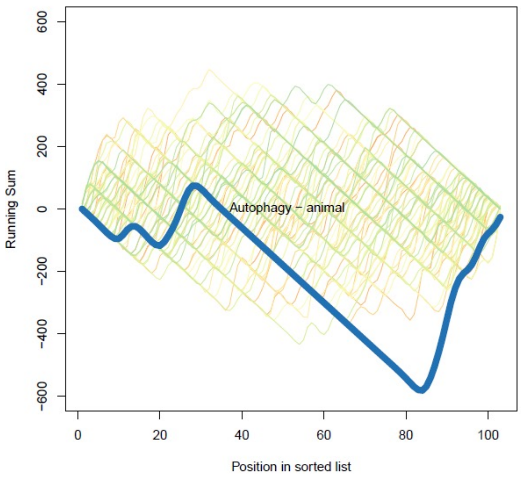


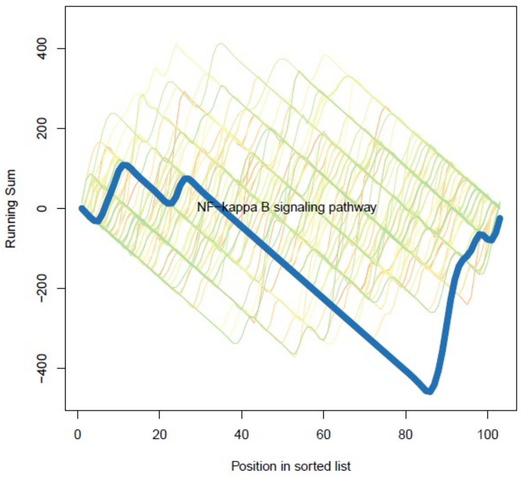


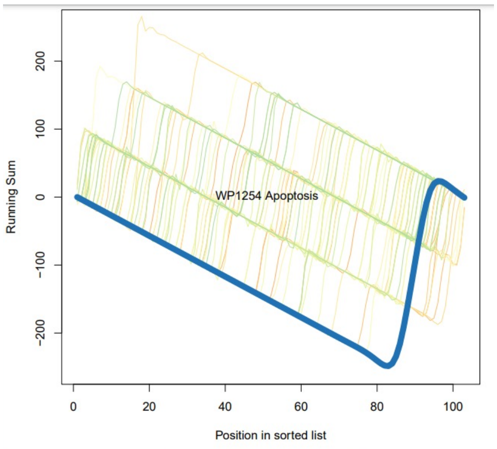

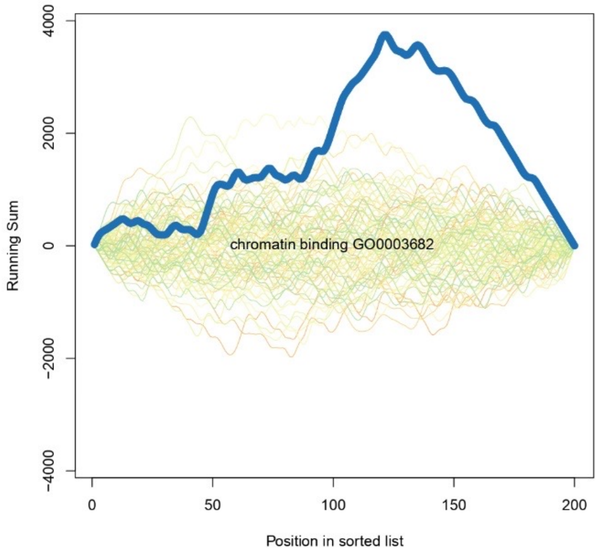

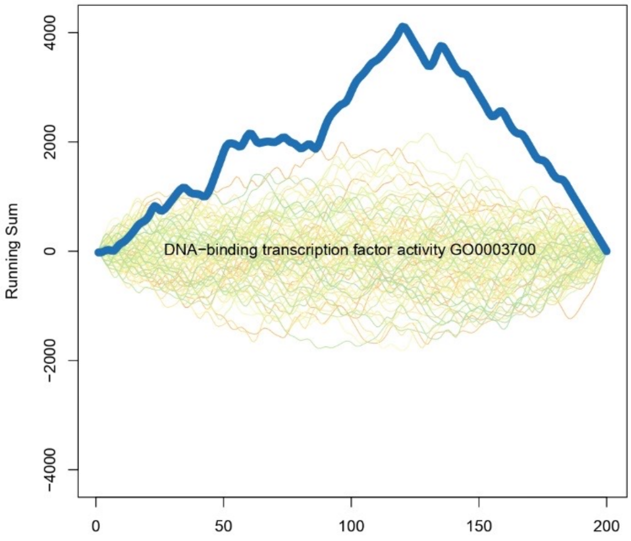

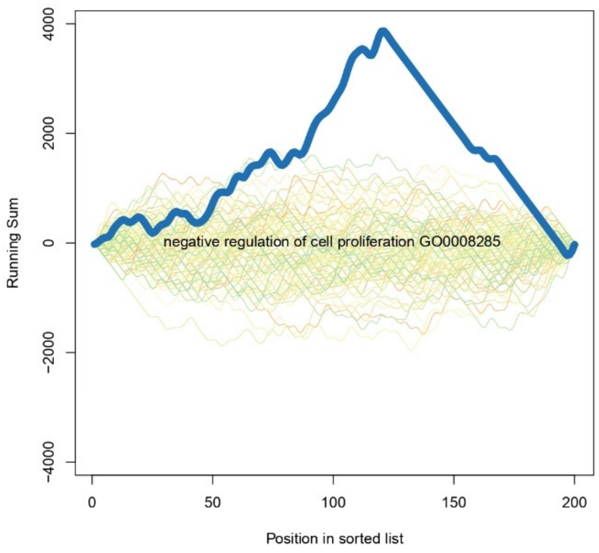

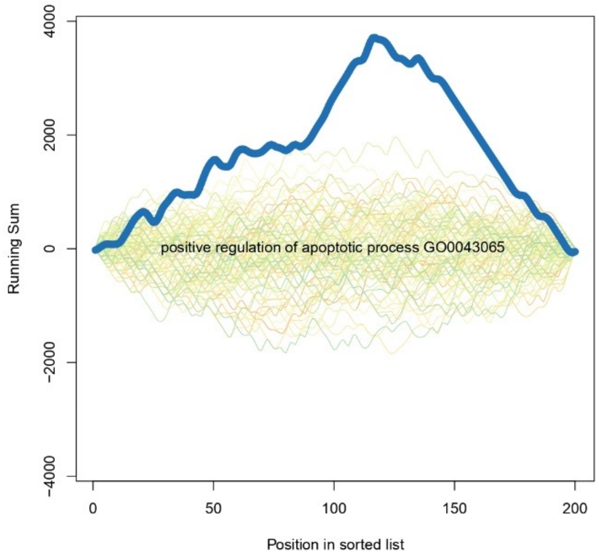

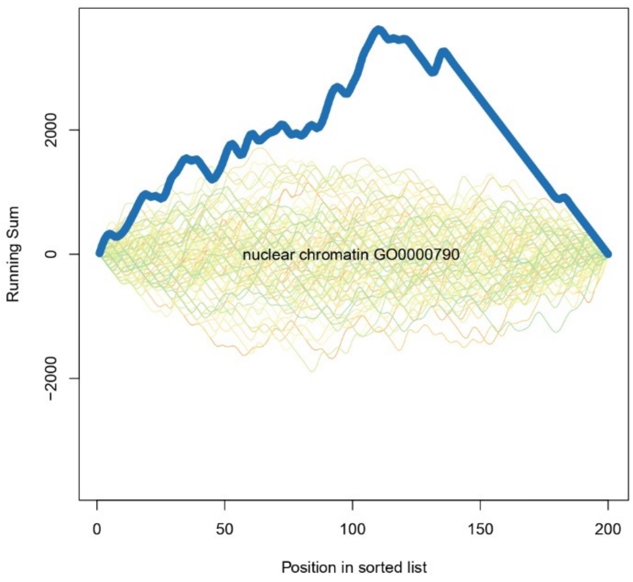

#### Transcription factor

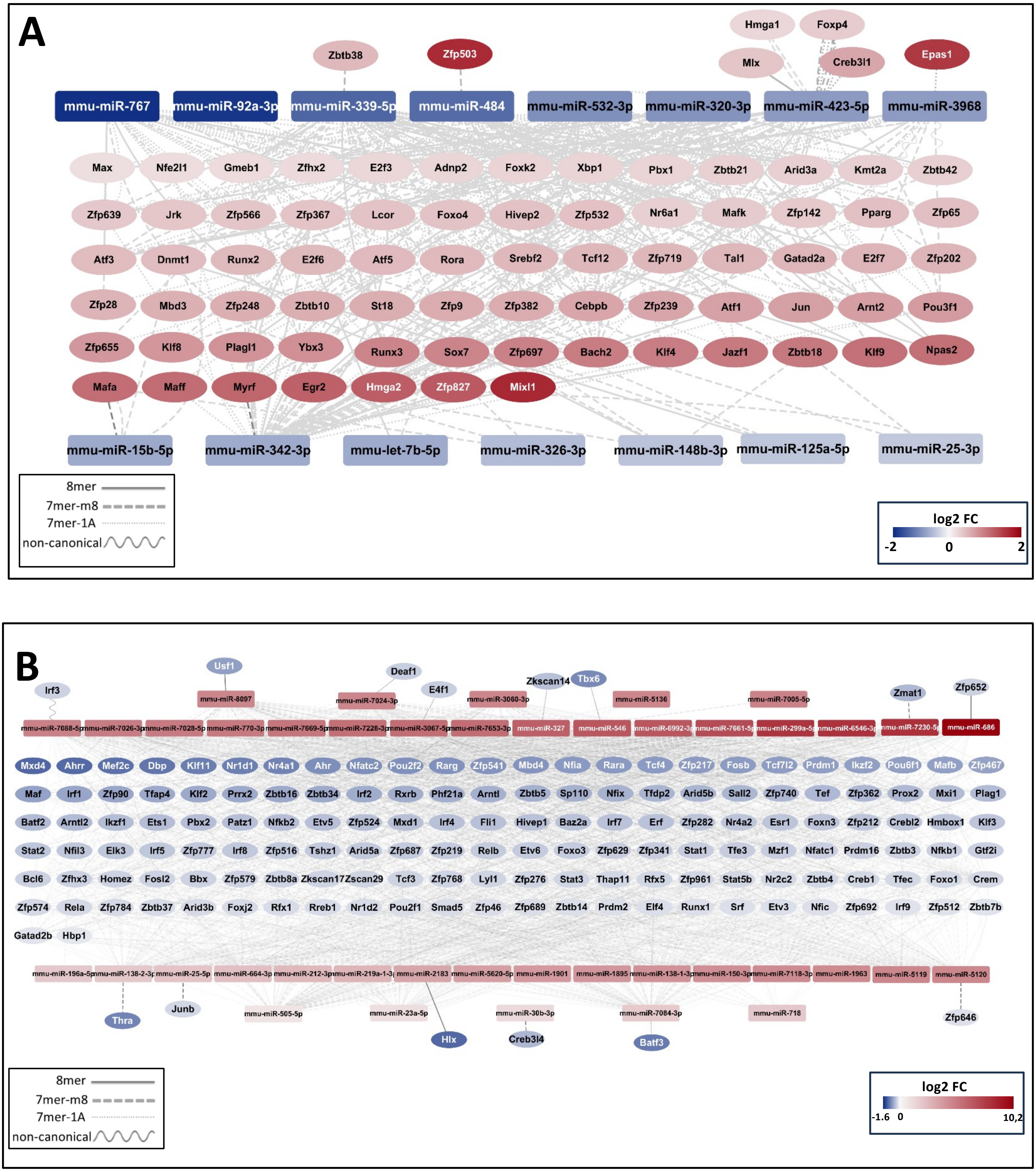

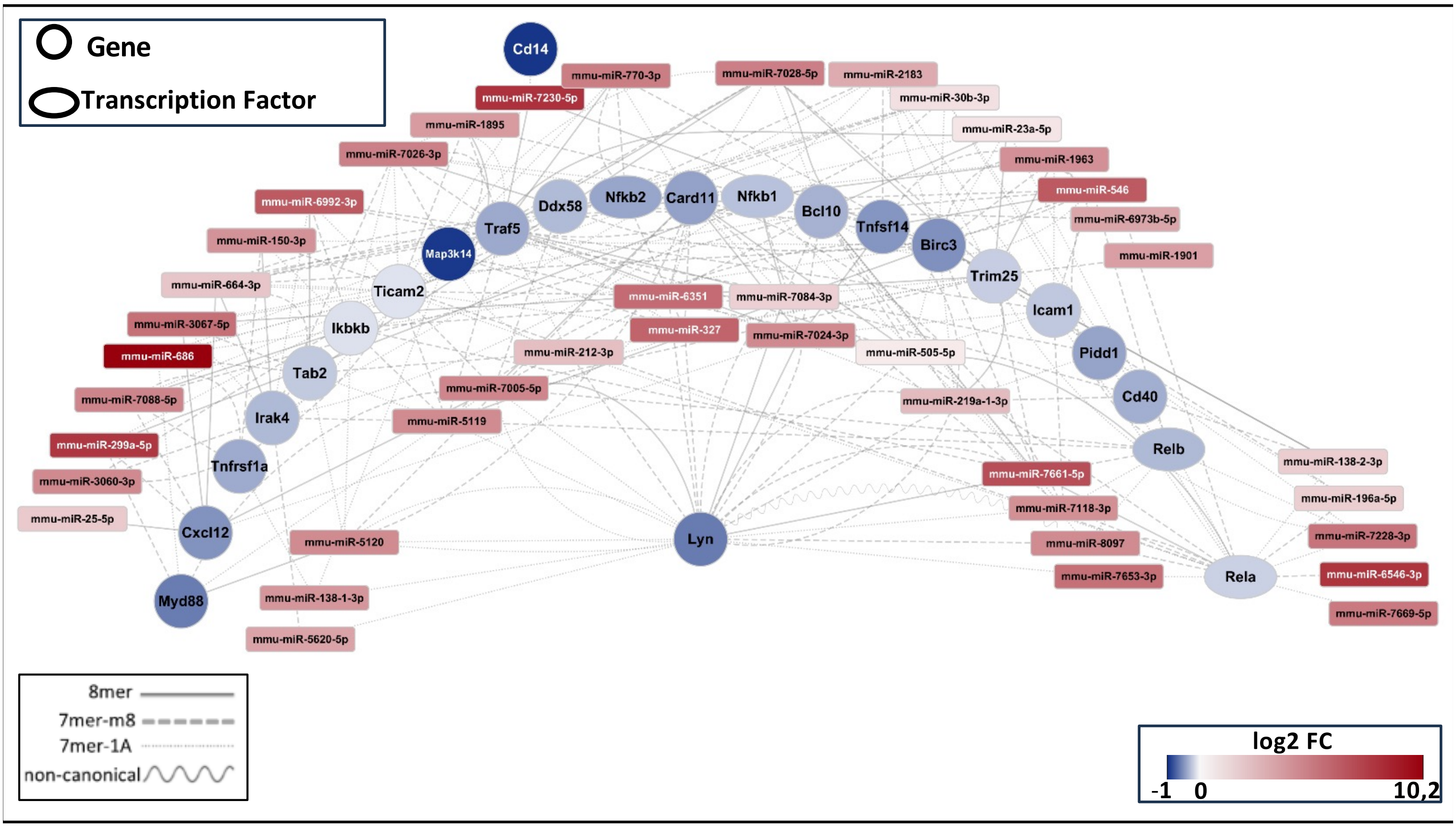

#### Cholesterol biosynthetic process.

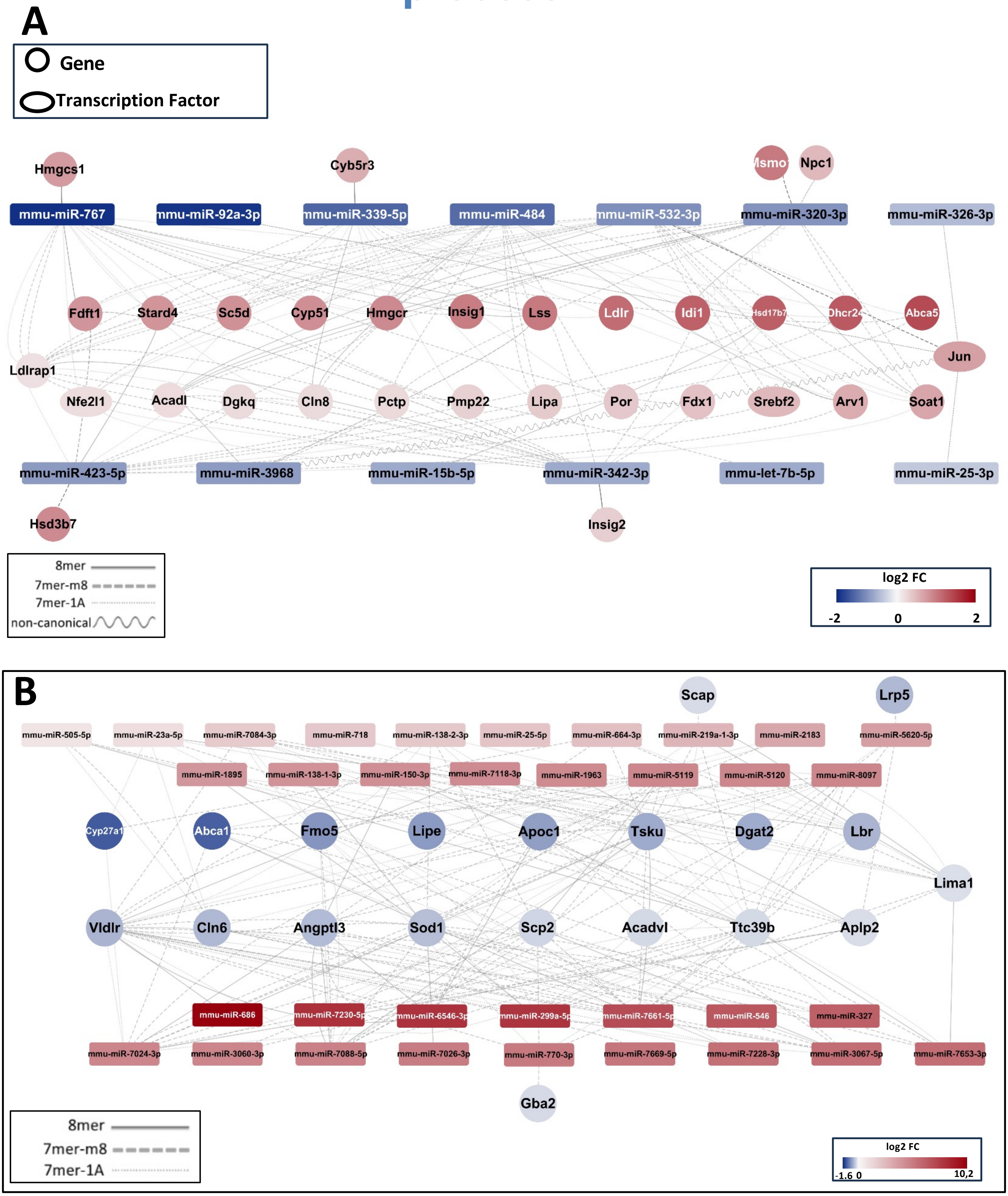
